# Patterns of NFkB activation resulting from damage, reactive microglia, cytokines, and growth factors in the mouse retina

**DOI:** 10.1101/2022.08.16.504164

**Authors:** Isabella Palazzo, Lisa Kelly, Lindsay Koenig, Andy J. Fischer

## Abstract

Müller glia are a cellular source for neuronal regeneration in vertebrate retinas. However, the capacity for retinal regeneration varies widely across species. Understanding the mechanisms that regulate the reprogramming of Müller glia into progenitor cells is key to reversing the loss of vision that occurs with retinal diseases. In the mammalian retina, NFκB signaling promotes glial reactivity and represses the reprogramming of Müller glia into progenitor cells. Here we investigate different cytokines, growth factors, cell signaling pathways, and damage paradigms that influence NFκB-signaling in the mouse retina. We find that exogenous TNF and IL1β potently activate NFκB-signaling in Müller glia in undamaged retinas, and this activation is independent of microglia. By comparison, TLR1/2 agonist indirectly activates NFκB-signaling in Müller glia, and this activation depends on the presence of microglia as *Tlr2* is predominantly expressed by microglia, but not other types of retinal cells. Exogenous FGF2 did not activate NFκB-signaling, whereas CNTF, Osteopontin, WNT4, or inhibition of GSK3β activated NFκB in Müller glia in the absence of neuronal damage. By comparison, dexamethasone, a glucocorticoid agonist, suppressed NFκB-signaling in Müller glia in damaged retinas, in addition to reducing numbers of dying cells and the accumulation of reactive microglia. Although NMDA-induced retinal damage activated NFκB in Müller glia, optic nerve crush had no effect on NFκB activation within the retina, whereas glial cells within the optic nerve were responsive. We conclude that the NFκB pathway is activated in retinal Müller glia in response to many different cell signaling pathways, and activation often depends on signals produced by reactive microglia.

## Introduction

There is a growing body of work describing the various cell signaling pathways that regulate the reprogramming of Müller glia (MG) into MG-derived progenitor cells (MGPCs). Jak/Stat-, MAPK-, mTOR-, and WNT-signaling, as well as secreted factors from reactive microglia, have been shown to play crucial roles in regulating MG reprogramming in different vertebrate species (Fischer et al., 2009; Fischer et al., 2014b; Nelson et al., 2012; Ramachandran et al., 2011; Todd et al., 2016; Todd et al., 2020; Wan et al., 2014; White et al., 2017). We have recently shown that NFκB is a part of the regulatory network that drives reactive MG to restore quiescence and suppress gene modules that promote formation of neurogenic progenitors in mouse retina (Hoang et al., 2020; Palazzo et al., 2022). We found that NFκB-signaling in damaged retinas is manifested in MG only when reactive microglia are present. Further, we show that inhibition of NFκB-signaling results in diminished recruitment of immune cells in damaged retinas, increased neuronal survival, and increased formation of neuron-like cells from Ascl1-overexpressing MG (Palazzo et al., 2022).

Similar to the results seen in the mouse retina, NFκB-signaling in the chick retina suppresses the formation of proliferating MGPCs and this process depends on signals from reactive microglia (Palazzo et al., 2020). NFκB components are not prevalently expressed in MG in damaged zebrafish retinas wherein MGPCs effectively regenerate retinal neurons, unlike the MG in mouse retina where NFκB is rapidly upregulated (Hoang et al., 2020; Palazzo et al., 2022). Thus, it is likely that NFκB signaling is a key point in the divergence of regenerative capacity of MG across different species. Accordingly, the purpose of this study was to better understand the growth factors, cytokines, and cell signaling pathways that influence NFκB-signaling in the retina. Herein, we investigate crosstalk between NFκB signaling and cell-signaling pathways known to influence MG, and whether reactive microglia are needed to mediate these responses. We find that IL1β, TNF, and Osteopontin produced by reactive microglia/macrophage, as well as TLR-, CNTF- and WNT-signaling pathways all converge on NFκB signaling in MG in the mouse retina. Considering that NFκB-signaling represses reprogramming of mouse MG by promoting reactivity networks and anti-neurogenic factors, while increasing cell death in damage retinas (Palazzo et al., 2022), it is important to understand how different cell signaling pathways coordinate with NFκB to develop techniques to enhance neuronal survival and retinal regeneration.

## Methods

### Animals

The use of animals in these experiments was in accordance with the guidelines established by the National Institutes of Health and the IACUC committee at the Ohio State University. Mice were kept on a cycle of 12 hours light, 12 hours dark (lights on at 8:00 AM). Experiments were performed using NFκB-eGFP reporter, which mice have eGFP-expression driven by a chimeric promoter containing three HIV-NFκB cis elements (Magness et al., 2004).

### Oral administration of PLX5622

Mice were fed chow formulated with PLX5622 (1200 ppm; Plexxikon; D11100404i) *ad libitum* for a minimum of two weeks before experiments, and this diet was continued through the duration of each experiment.

### Injections

Mice were anesthetized via inhalation of 2.5% isoflurane in oxygen and intraocular injections performed as described previously (Todd and Fischer, 2015). For all experiments, the vitreous chamber of right eyes of mice were injected with the experimental compound and the contralateral left eyes were injected with a control vehicle. Compounds used in these studies are as follows: N-methyl-D-aspartate (NMDA; 1.5ul of 100mM in PBS; Sigma; M3262), recombinant mouse IL1β (200 ng/dose; R&D systems; 401-ML) recombinant mouse TNF (250 ng/dose; BioLegend; 575204), recombinant mouse Osteopontin (OPN; 200ng/dose; BioLegend; 763604), CNTF (300ng/dose; R&D systems; 557-NT), FGF2 (250 ng/dose; R&D systems; 233-FB), Dexamethasone (Dex; 250ng/dose; MP Biomedicals; 194561), WNT4 (1ug/dose; Abcam; ab236179); CU-T12-9 (TLR1/2 agonist; 2.5ug/dose; Tocris; 5414), CU-CPT22 (TLR antagonist; 2.5ug/dose; Tocris; 4884), and a cocktail of three different GSK3β inhibitors (referred to as ABC) which included 1-Azakenpaullone (1-AP; 500 ng/dose; Selleck Chemicals; S7193), BIO (500 ng/dose; R&D Systems; 3194), CHIR 99021 (500 ng/dose; R&D Systems; 4423).

### Optic Nerve Crush

Optic nerve crush (ONC) was performed as previously described (Templeton and Geisert, 2012). Dumont #N7 cross-action forceps were used to retro-orbitally compress the optic nerve for 10 s. Optic nerves were harvested 8 days post-ONC, fixed for 30 minutes in 4% PFA, processed for immunohistochemistry as described below, and whole-mounted optic nerves were examined using confocal microscopy.

### Fixation, sectioning, and immunocytochemistry

Retinas were fixed, sectioned, and immunolabeled as described previously (Fischer et al., 2008; Fischer et al., 2014a; Gallina et al., 2015). None of the observed labeling was due to non-specific labeling of secondary antibodies or auto-fluorescence because sections labeled with secondary antibodies alone were devoid of fluorescence. Primary antibodies used in this study are described in **Table 1**. Secondary antibodies included donkey-anti-goat-Alexa488/568 (Life Technologies A3214; A3214), goat-anti-rabbit-Alexa488/568 (Life Technologies A32731; A-11036); and goat-anti-mouse-Alexa488/568/647 (Life Technologies A32723; A-11004; A32728) diluted to 1:1000 in PBS plus 0.2% Triton X-100.

**Table 1:** Antibody table

| **Antibody** | **Dilution** | **Host** | **Clone/Catalog number** | **Source** |
| --- | --- | --- | --- | --- |
| Sox2 | 1:1000 | Goat | KOY0418121 | R&D Systems |
| Sox9 | 1:2000 | Rabbit | AB5535 | Millipore |
| Iba1 | 1:300 | Rabbit | 019-19741 | Wako Industries |
| Glutamine synthetase | 1:1000 | Mouse | 610517 | BD Biosciences |
| pERK1/2 | 1:800 | Rabbit | 137F5 | Cell Signaling |
| pS6 | 1:750 | Rabbit | 2215 | Cell Signaling |
| pStat3 | 1:100 | Rabbit | 9131 | Cell Signaling |
| S100-beta | 1:500 | Rabbit | 37A | Swant<br>Immunochemicals |
| GFP | 1:1000 | Chicken | Ab13970 | Abcam |

**Table 2.** Pharmacological Compounds

| **Drug Name** | **Dose** | **Source** | **Catalog number** |
| --- | --- | --- | --- |
| NMDA | 15ug | Sigma | M3262 |
| Recombinant mouse IL1b | 200ng | R&D Systems | 401-ML |
| Recombinant mouse TNF | 250ng | BioLegend | 575204 |
| Recombinant mouse Osteopontin | 200ng | BioLegend | 763604 |
| FGF2 | 250ng | R&D Systems | 233-FB |
| Dexamethasone | 250ng | MP Biomedicals | 194561 |
| WNT4 | 1ug | Abcam | ab236179 |
| CU-T12-9 | 2.5ug | Tocris | 5414 |
| CU-CPT22 | 2.5ug | Tocris | 4884 |
| 1-Azakenpaullone | 500ng | Selleck Chemicals | S7193 |
| BIO | 500ng | R&D Systems | 3194 |
| CHIR 99021 | 500ng | R&D Systems | 4423 |

### Photography, immunofluorescence measurements, and statistics

Wide field photomicroscopy was performed using a Leica DM5000B microscope equipped with epifluorescence and Leica DC500 digital camera or Zeiss AxioImager M2 equipped with epifluorescence and Zeiss AxioCam MRc. Confocal images were obtained using a Leica SP8 imaging system at the Department of Neuroscience Imaging Facility at the Ohio State University. Images were optimized for color, brightness and contrast, multiple channels over laid and figures constructed using Adobe Photoshop. Cell counts were performed on representative images. To avoid the possibility of region-specific differences within the retina, cell counts were consistently made from the same region of retina for each data set.

Where significance of difference was determined between two treatment groups accounting for inter-individual variability (means of treated-control values) we performed a two-tailed, paired t-test. Where significance of difference was determined between two treatment groups, we performed a two-tailed, unpaired t-test. Significance of difference between multiple groups was determined using ANOVA followed by Tukey’s test.

GraphPad Prism 6 was used for statistical analyses and generation of histograms and bar graphs.

### scRNA-seq

scRNA-seq libraries from Hoang et al. 2020 were analyzed. In short, retinas were acutely dissociated via papain digestion (Worthington Biochemicals). Dissociated cells were loaded onto the 10X Chromium Cell Controller with Chromium 3’ V2 reagents. Sequencing and library preparation was performed as previously described (Hoang et al., 2020). Gene-Cell matrices for scRNA-seq data for libraries from saline and NMDA-treated retinas are available through GitHub: https://github.com/jiewwwang/Singlecell-retinal-regeneration

Using Seurat toolkits (Powers and Satija, 2015; Satija et al., 2015), Uniform Manifold Approximation and Projection (UMAP) for dimensional reduction plots were generated from 9 separate cDNA libraries, including 2 replicates of control undamaged retinas, and retinas at different times after NMDA-treatment. Seurat was used to construct gene lists for differentially expressed genes (DEGs), violin/scatter plots, and dot plots. Significance of difference in violin/scatter plots was determined using a Wilcoxon Rank Sum test with Bonferroni correction. Genes that were used to identify different types of retinal cells included the following: (1) Müller glia: *Glul, Nes, Vim, Scl1a3, Rlbp1*, (2) microglia: *C1qa, C1qb, Csf1r, Apoe, Aif1* (3) ganglion cells: *Thy1, Pou4f2, Rbpms2, Nefl, Nefm,* (4) amacrine cells: *Gad67, Calb2, Tfap2a,* (5) horizontal cells: *Prox1, Calb2,* (6) bipolar cells: *Vsx1, Otx2, Grik1, Gabra1,* and (7) cone photoreceptors: *Gnat2, Opn1lw*, and (8) rod photoreceptors: *Rho, Nr2e3, Arr1*.

Activated and +activated MG were identified based on elevated expression levels of *Nes, Vim, Il6, Egr1, Atf3,* and *Tgfb2* as described previously (Campbell et al., 2021). The “+activated MG” cluster is primarily comprised of MG from retinas 3hr after NMDA-treatment, the “activated MG” cluster is comprised of MG from 6hr after NMDA-treatment, and the “return to resting MG” cluster is comprised of MG primarily from 12, 24, 36, 48 and 72hr after NMDA-treatment (see Fig. 3c).

## Results

### Microglia-derived signals are sufficient to activate NFκB-signaling in Müller glia

The NFκB pathway is a master regulator of pro-inflammatory cell signaling and is activated by various cytokines, including members of interleukin (IL) and tumor necrosis factor (TNF) families (Osborn et al., 1989). The primary source of pro-inflammatory cytokines are microglia, the resident immune cell of the retina. We have previously shown that NMDA-damage results in NFκB activation in MG in the mouse retina, and the ablation of microglia prior to damage prevents this activation (Palazzo et al., 2022). These findings suggest that factors secreted by reactive microglia activate NFκB-signaling in MG. After excitotoxic retinal injury, microglia, but not other retinal cell types, rapidly upregulate TNF and IL1β (Todd et al., 2019) (**Fig. 1a-d)**, thus we tested whether IL1β and TNF activate NFκB-signaling in the retina in the absence of damage. We injected recombinant mouse IL1β intravitreally in the eyes of NFκB-eGFP reporter mice, which have eGFP-expression driven by a chimeric promoter containing three HIV-NFκB cis elements (Magness et al., 2004). We harvested retinas 24h post-injection and found a robust induction of the NFκB-eGFP reporter in MG from retinas treated with a single intravitreal injection of IL1β **(Fig. 1e,f)**. MG were unambiguously identified based on their distinct morphology within the retina (Cajal, 1972). Next, we sought to determine if IL1 β induced the NFκB reporter in the absence of microglia. We fed mice a diet with PLX5622, a Csf1r antagonist that results in ablation of microglia throughout the CNS (Elmore et al., 2014). We found that treatment with exogenous IL1β induced NFκB activation in MG in the absence of microglia and retinal damage (**Fig. 1e,f**). We confirmed that microglia were ablated by labeling for Iba1 (**Fig. 1f**). We have previously reported rapid and robust upregulation of this NFκB-eGFP reporter following NMDA damage (Palazzo et al., 2022) and confirm this expression 48h after damage **(Fig. 1g)**. Our previous study indicated that damage-induced activation of NFκB signaling is greatly diminished when microglia are ablated (Palazzo et al., 2022). Thus, we tested whether this deficit in NFκB activation in damaged retinas could be rescued by exogenous IL1 β treatment. The PLX5622 diet was maintained for the duration of the experiment, and intravitreal injections of NMDA alone or with IL1β were performed 2 weeks after the onset of the diet. Retinas were harvested 24hrs after injection.

**Figure 1:**
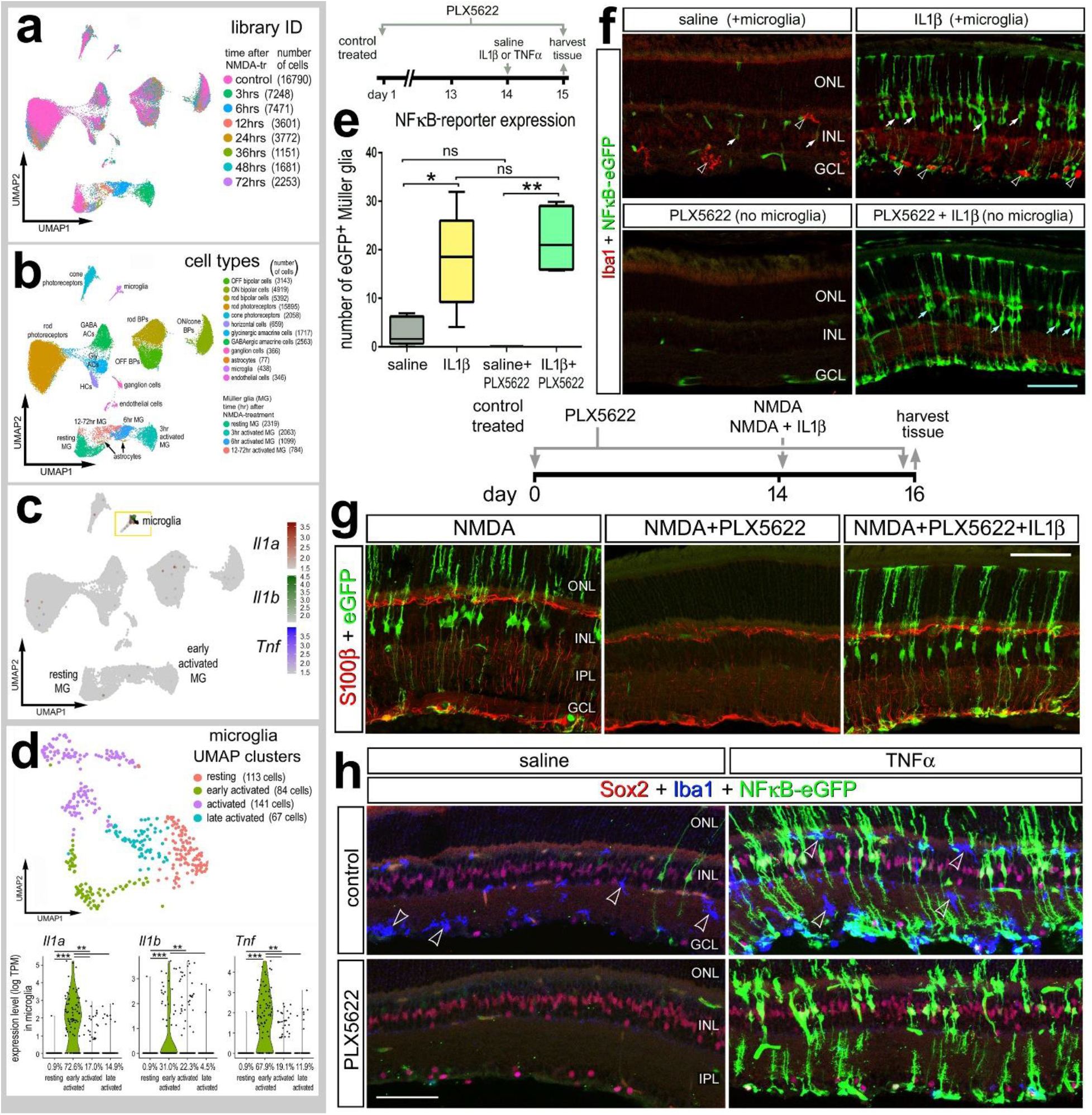
Exogenous cytokines stimulate NFκB-signaling in MG independent of microglia. Aggregated scRNA-seq libraries were prepared from WT control retinas and retinas 3h, 6h,12h, 24h, 36h, 48h, and 72h after NMDA damage **(a)**. Clusters were identified based on expression of cell-distinguishing markers described in Methods **(b)**. UMAP feature plot illustrates expression of Il1a, Il1b and Tnf across all retinal cells **(c)** and microglia **(d)**. NFκB-eGFP mice were fed a control diet or diet containing PLX5622 for 2 weeks preceding a single intraocular injection of saline, NMDA, IL1β, or TNF. Retinas were harvested 24h after IL1β **(e,f)**, NMDA+IL1β **(g)**, or TNF **(h)**injection, and retinal sections were immunolabeled for GFP (green; **f-h**), Iba1(red; **f**) (blue; **h**), S100b (red; **g**), or Sox2 (red; **h**). **(e)** Boxplots illustrate mean, upper quartile, lower quartile, upper extreme, and lower extreme of total number of GFP^+^ MG (n=5 animals). Significance of difference (*p<0.05, **p<0.005) between treatment groups was determined via One-way ANOVA with Tukey’s test. Hollow arrows indicate microglia **(f)**, solid arrows indicate MG **(f)** or microglia **(h)**. Calibration bars panels **f** and **h** represent 50 μm. Abbreviations: ONL – outer nuclear layer, INL – inner nuclear layer, IPL – inner plexiform layer, GCL – ganglion cell layer.

Consistent with previous findings, we found the NFκB reporter prominently active in MG following NMDA-treatment, and this activation was absent when microglia were ablated (**Fig. 1g**). However, a single intraocular injection of IL1β concurrent with NMDA resulted in activation of the NFκB reporter in MG despite the absence of microglia (**Fig. 1g**).

We next investigated whether exogenous TNF is sufficient to activate NFκB signaling in MG in the absence of damage and microglia. We found that a single intraocular injection of TNF robustly induced NFκB-eGFP reporter expression in MG in undamaged retinas, indicated by GFP^+^ cells expressing MG marker Sox2 (**Fig. 1h**). Additionally, we injected TNF following microglia ablation via PLX5622 and found widespread NFκB activity in MG independent of microglia (**Fig. 1h**). Taken together, our findings suggest that microglia derived IL1β and TNF directly activate NFκB-signaling in MG.

### Stimulation of TLR signaling activates NFκB-signaling in MG

Toll-like receptors (TLRs) are pattern recognition receptors for damage-associated molecular patterns (DAMPs) and pathogen-associated molecular patterns (PAMPs) that mediate both innate and adaptive immune responses. TLRs are responsive to DAMPs associated with cell death and tissue damage (reviewed by (Piccinini and Midwood, 2010). Activation of TLRs can induce expression of pro-inflammatory cytokines and chemokines, and in immune cells, TLR activation directly signals through NFκB via the IKK complex (reviewed by (Kawai and Akira, 2010)). Accordingly, we investigated whether signaling through TLRs influence NFκB signaling in the retina.

To establish a cellular context for TLR signaling in the retina, we queried single cell (sc) RNA-seq libraries from undamaged and NMDA-damaged retinas (Hoang et al., 2020) **(Fig. 2a-e)**. We found that *Tlr1, Tlr4,* and *Tlr7* were expressed at relatively low levels in relatively few microglia (**Fig. 2a-d**). By comparison, *Tlr2* was detected microglia and also detected in a few activated MG after damage (**Fig. 2a-d**). We bioinformatically isolated microglia and re-normalized to re-order these cells in a UMAP plot (**Fig. 2c-e**). We probed for expression of TLRs in the microglia and found that *Tlr2* is upregulated rapidly (3hrs after NMDA) in early activated microglia, whereas *Tlr1, Tlr4,* and *Tlr7*are expressed at low levels (**Fig. 2e**).

**Figure 2:**
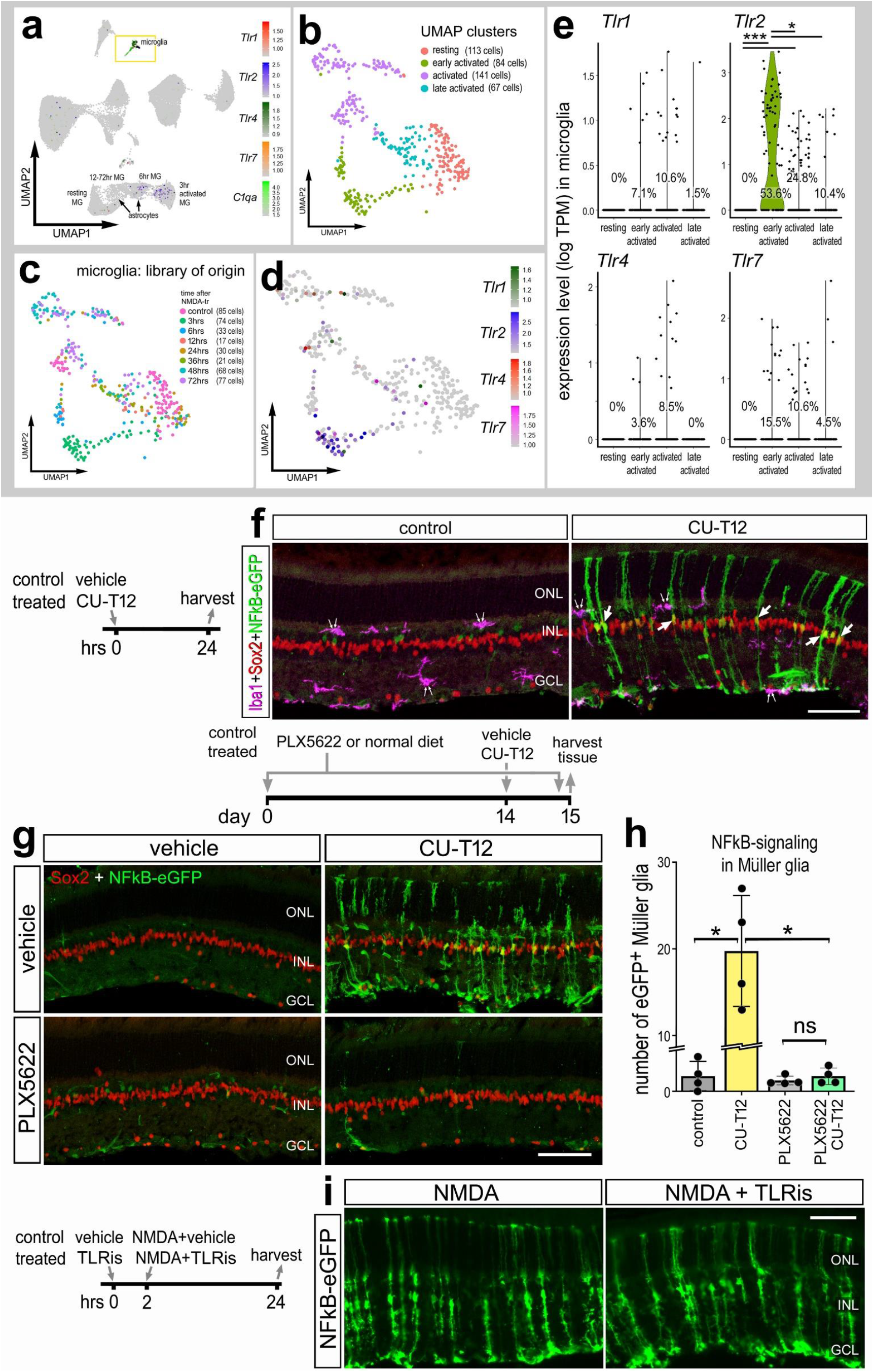
TLRs are expressed by microglia. Aggregated scRNA-seq libraries were prepared from WT control retinas and retinas 3h, 6h,12h, 24h, 36h, 48h, and 72h after NMDA damage. Clusters were identified based on expression of cell-distinguishing markers described in Methods. UMAP feature plot illustrates expression of *Tlr1, Tlr2, Tlr4, Tlr7,* and *C1qa* across all retinal cells **(a)**. UMAP plots of isolated and re-embedded microglia show distinct clusters **(b,c)**. UMAP plot **(d)** and violin plots **(e)** illustrate levels of expression of *Tlr1, Tlr2, Tlr4, and Tlr7* in microglia clusters. Each dot represents one cell. Significance of difference (**p<0.01, ***p<0.001) was determined via Wilcoxon rank sum with Bonferroni correction **(e)**. **(f)** Eyes of NFκB-eGFP mice were injected with TLR agonist (CU-T12) or vehicle (control), retinas were harvested 24hrs later, and retinal sections were immunolabeled for Iba1 (magenta; **f**), Sox2 (red; **f,g**), and GFP (green; **f,g,i**). Double arrows in **(f)** indicate microglia and single arrows indicate GFP^+^ MG. **(g)** NFκB-eGFP mice were fed a control diet or diet containing PLX5622 for 2 weeks prior to a single intravitreal injection of vehicle (control) or CU-T12, retinas were harvested 24 hrs later. **(h)** Histogram illustrates the mean ± SD (individual datapoints shown) number of GFP^+^ MG. Significance (*p<0.01) of difference was determined via one-way ANOVA followed by post-hoc Man-Whitney U-test. **(i)** The eyes of NFκB-eGFP mice were injected with vehicle or TLR antagonist (CU-CPT22; TLRi), followed 2hrs later by injection of NMDA+vehicle or NMDA+TLRi, and retinas harvested 24hrs after the last injection. Calibration bars represents 50 μm. Abbreviations: ONL – outer nuclear layer, INL – inner nuclear layer, IPL – inner plexiform layer, GCL – ganglion cell layer, NS – not significant.

**Figure 3:**
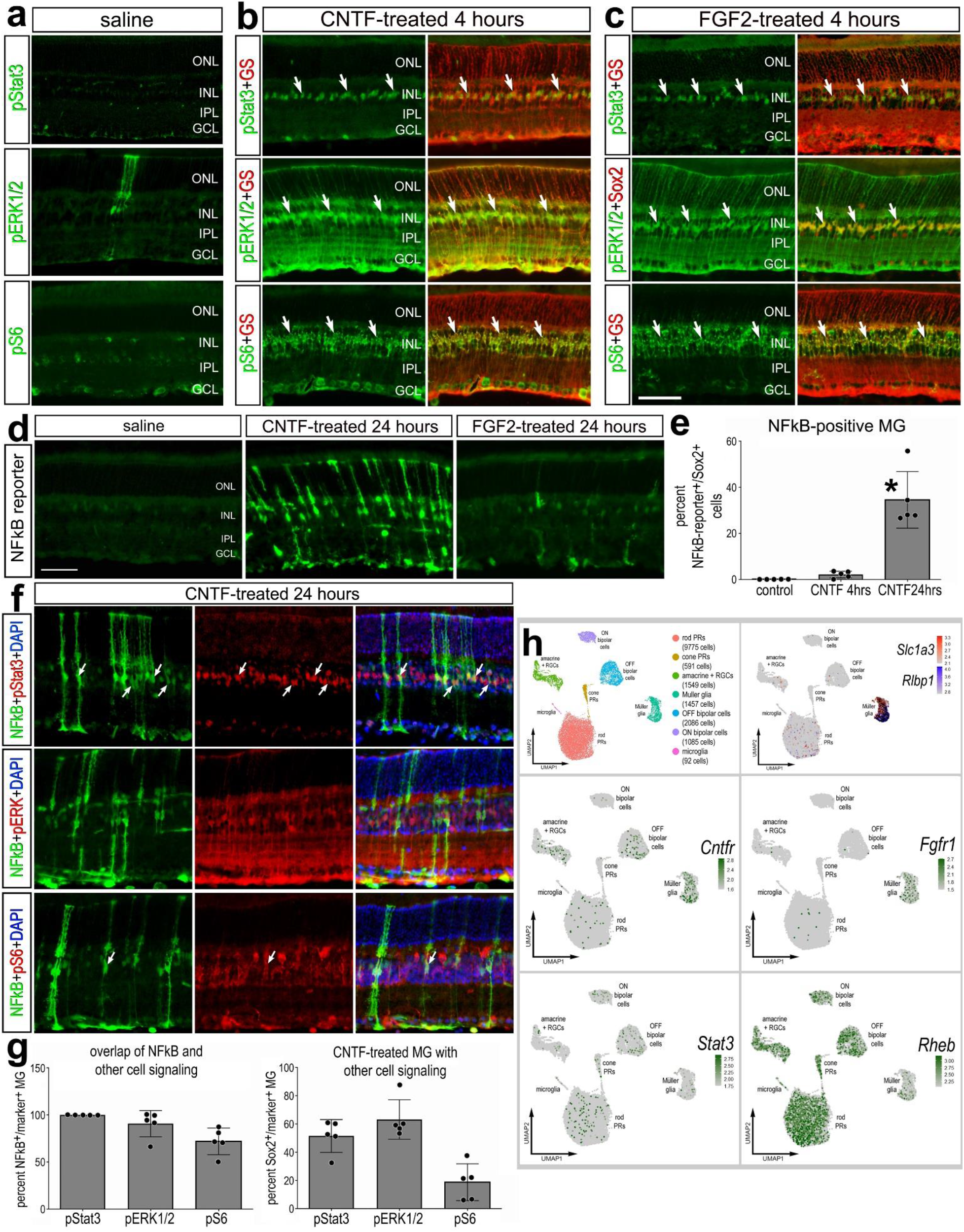
CNTF activates NFκB signaling; FGF2 does not. Eyes of NFκB-eGFP or control mice were injected with vehicle (saline), CNTF, or FGF2 and retinas were harvested 4hrs **(a-c)** or 24hrs later **(d-g)**. Retinal sections were immunolabeled with antibodies to GS (red – **a-c**) or Sox2 (red – **a-c**) and pStat3 (green – **b,c**; red -**f**), pERK1/2 (green – **b,c**; red -**f**), pS6 (green – **b,c**; red -**f**), or GFP (green – **d,f**). **(b)** scRNA-seq libraries were prepared from WT undamaged retinas. The histogram in panel **e** illustrates the mean (± SD and individual biological replicates) percentage of GFP^+^ MG over total MG (Sox2+). Significance (*p<0.01) of difference was determined via ANOVA followed by post-hoc Man-Whitney U-test. The histogram in panel **g** illustrates the mean (± SD and individual biological replicates) percentage of GFP^+^ MG that are co-labeled for pStat3, pERK1/2 or pS6 or the percentage of total MG that are labeled for pStat3, pERK1/2 or pS6 at 24 hours after CNTF-treatment. Arrows indicate double-labeled MG. Abbreviations: ONL – outer nuclear layer, INL – inner nuclear layer, IPL – inner plexiform layer, GCL – ganglion cell layer. **(h)** MG were identified based on expression of cell-distinguishing markers including *Slc1a3* and *Rlbp1,* and other types of retinal cells were identified as described in the Methods. **(h)** UMAP feature plots illustrate expression of *Cntfr, Stat3, Fgfr1*, and *Rheb* across all retinal cells.

We next treated retinas of NFκB-eGFP reporter mice with a small molecule agonist (CU-T12), which has been shown to selectively activate TLR1 and TLR2 (Cheng et al., 2015). We found that a single intravitreal injection of CU-T12 induced NFκB activity in Sox2^+^ MG, but not in Iba1 ^+^ microglia (**Fig. 2f**). These data indicate that activation of TLR1/2 is sufficient to induce NFκB activity in MG in the absence of retinal damage. We next tested whether TLR1/2 agonist acts directly on MG or indirectly via microglia. Accordingly, we ablated microglia via PLX5622 treatment prior to a single intravitreal injection of TLR1/2 agonist, CU-T12, and harvested retinas 24hrs postinjection. We found a significant induction of the NFκB-eGFP reporter in MG when microglia were present, however treatment with TLR1/2 agonist failed to activate NFκB in MG when the microglia were absent (**Fig. 2g-h**). This finding suggests that TLR1/2 agonist likely acts through microglia to produce signals that secondarily induce activation of NFκB-signaling in MG. By comparison, we asked if inhibition of TLRs suppressed NFκB-signaling in MG. We treated NMDA damaged retinas with CU-CPT22, a small molecule antagonists of TLR1/2 (Cheng et al., 2012). We found that inhibition of TLRs did not suppress NMDA-damage induced activation of NFκB in MG (**Fig. 2i**). This suggests that inhibition of TLRs, expressed primarily by microglia, is not sufficient to suppress the production of factors that activate NFκB-signaling in MG.

### CNTF activates NFκB-signaling; FGF2 does not

In the zebrafish retina, ciliary neurotrophic factor (CNTF) treatment stimulates the proliferation of MGPCs in the absence of damage by signaling though Janus kinase signal transducers and activators of transcription (Jak/Stat) (Nelson et al., 2012; Zhao et al., 2014). By comparison, Jak/Stat signaling suppresses the formation of neuron-like cells from Ascl1-overexpressing MG in rodent retinas (Jorstad et al., 2020) and promotes MGPC formation and proliferation, but suppresses neuronal differentiation from MGPCs in chick retinas (Todd et al., 2016). In immune cells, Jak/Stat signaling is associated with inflammatory responses that may coordinate with NFκB (Frühbeck, 2006). Accordingly, we probed for crosstalk between Jak/Stat- and NFκB-signaling.

CNTF is known to potently activate Jak/Stat signaling in the retina (Todd et al., 2016; Wen et al., 2012). At 4 and 24 hours after injection, we found that CNTF induced upregulation of Jak/Stat-, MAPK-, and mTor-active cell signaling markers, indicated by increased expression of phospho-Stat3 (pStat3), phospho-ERK1/2 (pERK1/2), and phospho-S6 (pS6) in MG that express glutamine synthetase (GS) or Sox2 (**Fig. 3a-c**). We found that CNTF treatment resulted in a robust activation of the NFκB-reporter in more than one third of the MG with at 24 hours, but not 4 hours, after treatment (**Fig. 3d,e**). More than 94% of the MG were positive for pStat3, pERK1/2 or pS6 at 4 hours after CNTF treatment. We next assessed the potential crosstalk between FGF/MAPK and NFκB-signaling. MAPK-signaling is known to promote the formation of MGPC in fish, chick, and mouse retina (Fischer et al., 2009; Karl et al., 2008; Wan et al., 2014). Unlike CNTF-treatment, FGF2-treatment had little effect upon the activated of NFκB-reporter, whereas there was a robust accumulation of pStat3, pERK1/2 and pS6 in MG at 4 or 24 hours after treatment (**Fig. 3d**). We next investigated whether there was overlap of NFκB with pStat3, pERK1/2, and pS6 at 24 hours after CNTF-treatment.

100% of the NFκB+ MG were positive for pStat3, >90% were positive for pERK1/2, and >70% were positive for pS6 (**Fig. 3g**). At 24 hours after CNTF-treatment, the percentage of MG that were positive pStat3, pERK1/2 and pS6 was reduced to about 50%, 60% and 20%, respectively (**Fig. 3g**). Taken together, these data indicate that NFκB is activated by CNTF, but not FGF2, in a delayed manner and may act in parallel and coordinate with pStat3, mTOR, and MAPK signaling.

To better understand the cellular context for CNTF- and FGF2-mediated activation of Stat- and MAPK-signaling in MG we probed scRNA-seq libraries for cell signaling components and receptors in WT undamaged retinas **(Fig. 3h)**. Clusters of MG were identified based on expression of *Slc1a3* and *Rlbp1* **(Fig. 3h)**. Consistent with previous reports (Fuhrmann et al., 1998), *Cntfr* was detected in some inner retinal neurons (amacrine and bipolar cells) as well as resting MG (**Fig. 3h)**, and *Fgfr1* was primarily expressed by resting MG **(Fig. 3h**). *Rheb* (directly upstream of mTorc1) and *Stat3* are widely expressed by most retinal cell types (**Fig. 3h**). These findings suggest that treatment of retinal cells with CNTF or FGF2 results in activation of cell signaling that is prominent in MG, and this pattern of activation likely results from restricted expression of *Cntfr* and *Fgfr1* in MG.

### WNT signaling activates NFκB-signaling

Wnt/β-catenin-signaling is a key signaling pathway that regulates MG reprogramming (Das et al., 2006; Gallina et al., 2015; Meyers et al., 2012; Osakada et al., 2007; Ramachandran et al., 2011; Yao et al., 2016). There is evidence of bidirectional communication between Wnt signaling and NFκB signaling (reviewed by (Ma and Hottiger, 2016). Since NFκB suppresses neurogenic programs in MG (Palazzo et al., 2020; Palazzo et al., 2022), we probed for cross-talk between WNT and NFκB signaling.

We injected recombinant WNT4 ligand intravitreally in NFκB-eGFP mice and assessed reporter activity 24 hours after treatment. We found treatment with WNT4 was sufficient to induce NFκB activity in MG (**Fig. 4a,b**). Wnt-signaling is activated by Wnt ligands binding to receptors and subsequent inactivation GSK3β, an enzyme that mediates phosphorylation of β-catenin to target for degradation. Thus, upon inactivation of GSK3β, β-catenin accumulates and translocates to the nucleus to activate transcription of target genes (Behrens et al., 1996). Previous studies indicate that GSK3β can regulate NFκB-signaling independent of Wnt/β-catenin (Saegusa et al., 2007). Thus, we next targeted GSK3β directly. We treated retinas with a cocktail of 3 different GSK3β inhibitors, 1-AP, BIO and CHIR, (GSK3βi) as described in the Methods. This combination of small molecule inhibitors has previously been shown to effectively activate GSK3β-mediated WNT signaling in the retina (Gallina et al., 2015). We found that a single injection of the GSK3β inhibitors results in activation of NFκB in undamaged retinas (**Fig 4c,d**). Together, these data indicate that WNT4 is capable of stimulating NFκB activity in MG, and this may be mediated inactivation of GSK3β.

**Figure 4:**
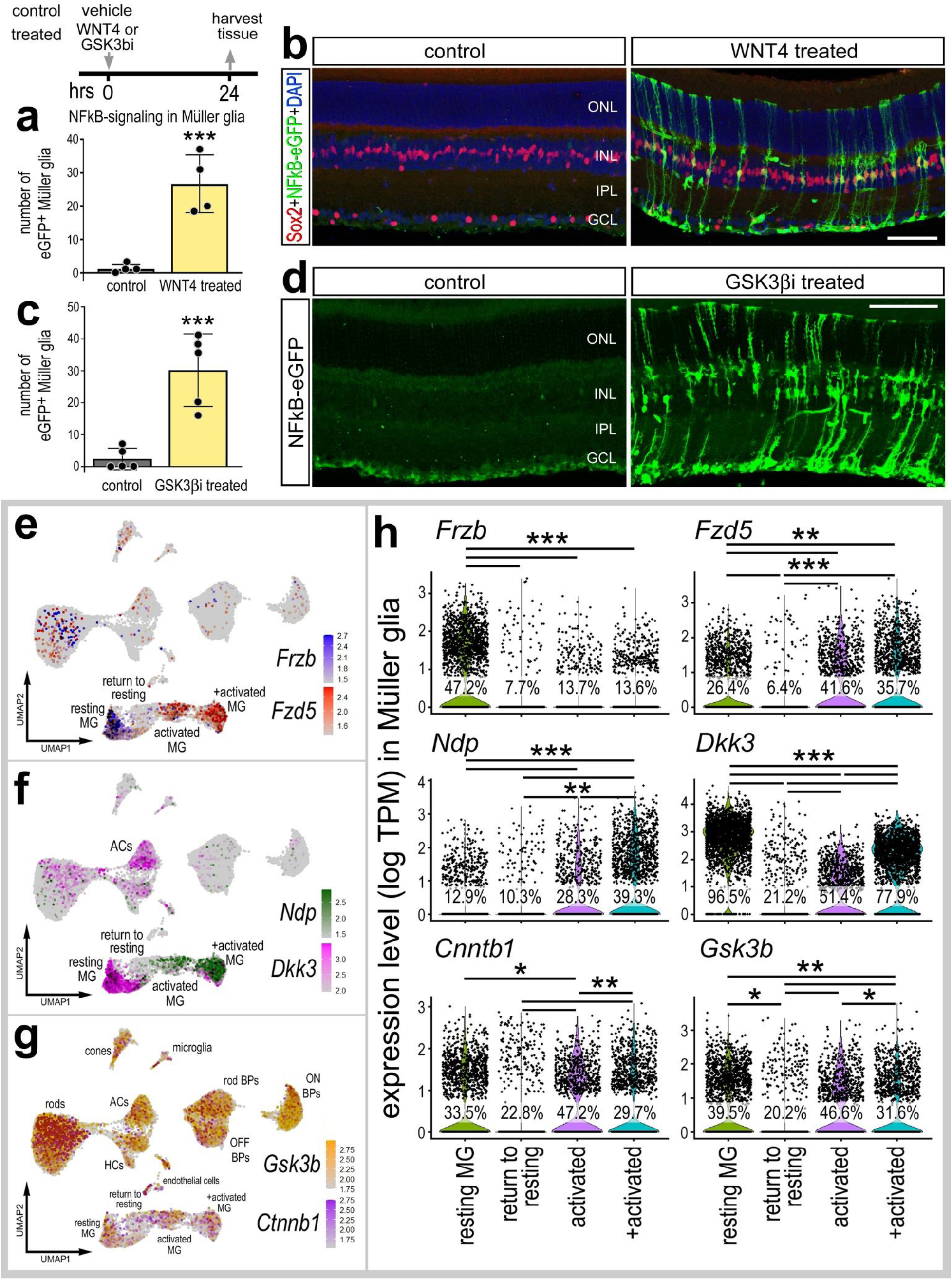
WNT4 or GSK3β-inhibitors activate NFκB. Eyes of NFκB-eGFP mice were injected with vehicle (control), WNT4, or a cocktail of GSK3β inhibitors described in Methods (GSK3βi), and retinas were harvested 24hrs post-injection. Retinal sections were immunolabeled with antibodies to Sox2 (red; **b**) and GFP (green; **b,d**). Histograms illustrate the mean ± SD (with individual datapoints shown) number of GFP^+^ MG **(a,c)**. Significance (***p<0.001) of difference was determined via paired t-test. **(e-h)** Aggregated scRNA-seq libraries were prepared from WT control retinas and retinas 3h, 6h,12h, 24h, 36h, 48h, and 72h after NMDA damage. **(e-g)** UMAP feature plots illustrate patterns of expression of *Frzb, Fzd5, Ndp, Dkk3, Gsk3b* and *Ctnnb1* across all retinal cells. **(h)** Violin plots illustrate levels of expression of *Frzb, Fzd5, Ndp, Dkk3, Gsk3b* and *Ctnnb1* in isolated MG clusters. Each dot represents one cell. Significance (*p<0.01; **p<10^-10^, ***p<10^-20^) of difference was determined by using a Wilcoxon Rank Sum with Bonferroni correct. Calibration bars represent 50 μm. Abbreviations: ONL – outer nuclear layer, INL – inner nuclear layer, IPL – inner plexiform layer, GCL – ganglion cell layer.

To better understand the cellular context for activation of NFκB in MG following treatment with WNT4 and GSK3β-inhibitors we probed scRNA-seq libraries for WNT-related signaling components in undamaged and NMDA-damaged retinas. WNT4 binds to Frizzled receptors and LRP co-receptors (Dijksterhuis et al., 2015; Ring et al., 2014). We found that WNT receptor *Fzd5* was expressed by resting MG (**Fig. 4e)**. *Fzd5* was rapidly upregulated at 3-12hr (activated MG clusters) and then downregulated at 24-48hrs (return to resting MG cluster) after NMDA (**Fig. 4h**). Other Frizzled receptor isoforms *(Fzd1, Fzd2, Fzd4, Fzd6)* were not highly expressed in our scRNA-seq libraries (not shown). We next probed for changes in expression of genes involved in regulating Wnt-signaling. *Ndp* facilitates the activation of Wnt-signaling (Seitz et al., 2010; Xu et al., 2004), whereas *Frzb* and *Dkk3* act to suppress Wnt-signaling by sequestering Wnt ligands (Lee et al., 2009; Veeck and Dahl, 2012). *Frzb* and *Dkk3* are highly expressed by resting MG and rapidly downregulated following damage (**Fig. 4f,h**), and is also detected in amacrine cells (**Fig. 4f**). By comparison, *Ndp* was expressed at low levels in resting MG, upregulated in activated MG, and was not expressed by other types of retinal cells (**Fig. 4f,h**). By contrast, *Gsk3b* and *Ctnnb1* (β-catenin) were widely expressed by most types of retinal cells (**Fig. 4g)**. Following NMDA-treatment, levels of *Gsk3b* and *Ctnnb1* are expressing in activated MG, but lower levels are detected in MG returning to a resting state (**Figs. 4g,h**). These findings indicate that *Frzb* is highly expressed by resting MG, but not other by other types of retinal cells. Thus, intraocular injections of Wnt likely act directly on MG. However, *Gsk3b* is widely expressed by many different types of retinal cells. Thus, the Gsk3b inhibitors may not be acting directly on MG to activate cell signaling. Collectively, these findings indicate that WNT ligands activate NFκB in MG independent of damage, and further that inhibition of GSK3β can activate NFκB-signaling independent of WNT activation.

### Glucocorticoid signaling suppresses NFκB-signaling

Activation of glucocorticoid signaling is potently anti-inflammatory in many different tissues and damage contexts (Ayroldi et al., 2012; Beck et al., 2009; Carrillo-de Sauvage et al., 2013). In the retina, glucocorticoids suppress inflammation and promote neuronal survival (Gallina, 2015; Iribarne and Hyde, 2021; Zhang et al., 2020). Additionally, glucocorticoid signaling in the chick retina suppresses the proliferation of MGPCs (Gallina et al., 2014). Accordingly, we probed for NFκB-eGFP reporter expression in NMDA-damaged retinas treated with dexamethasone, a glucocorticoid agonist. We found that dexamethasone treatment significantly reduced numbers of GFP^+^ MG in damaged retinas (**Fig. 5a-c**). In addition, dexamethasone significantly reduced the total number of Iba1^+^ of microglia/macrophage (**Fig. 5d-e**) and reduced numbers of dying cells in the INL, but not the GCL, in NMDA-damaged retinas (**Fig. 5f-i**). Collectively, these data suggest that activation of glucocorticoid signaling is neuroprotective, and this effect may, in part, be mediated by reduced NFκB-activation in MG and reduced accumulation of reactive microglia/macrophage.

**Figure 5:**
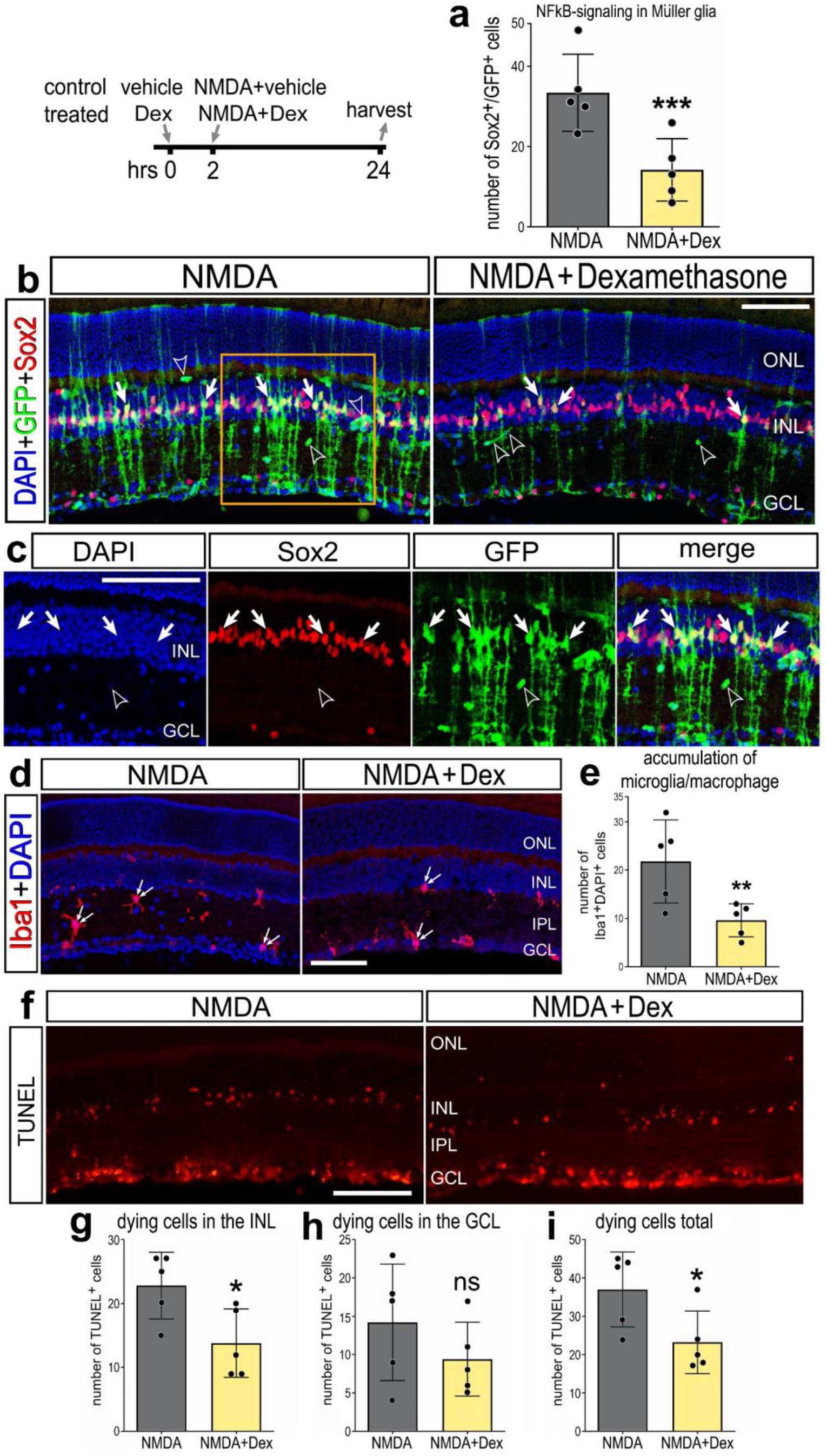
Dexamethasone suppresses NFκB activation in damaged retinas. Eyes of NFκB-eGFP mice were injected with dexamethasone or vehicle, followed 2hrs later by NMDA ± dexamethasone, and retinas harvested 24hrs post-injection. **(a)** Histogram illustrates mean ± SD (individual datapoints shown) number of GFP^+^ MG. **(b-d)** Retinal sections were immunolabeled with antibodies to Sox2 (red; **b,c**) and GFP (green; **b,c**), or Iba1 (red; **d**). **(c)** is enlarged region indicated in **(b)**. Retinal sections were labeled for TUNEL (red; **f**). Histograms illustrate the mean ± SD (individual datapoints shown) number of Iba1 ^+^cells **(e)** or number of TUNEL^+^ cells in the INL **(g**), GCL **(h)**, or total **(i)**. Significance (**p<0.01, ***p<0.001) of difference was determined via paired t-test. Solid arrows indicate GFP^+^ MG **(b)**, hollow arrowheads indicate blood vessels **(b)**, and small double-arrows indicate microglia/macrophage **(d)**. Calibration bars represent 50 μm. Abbreviations: ONL – outer nuclear layer, INL – inner nuclear layer, IPL – inner plexiform layer, GCL – ganglion cell layer.

### Osteopontin is expressed by microglia and activates NFκB-signaling in MG

In addition to other pro-inflammatory cytokines, reactive microglia upregulate osteopontin (OPN) in degenerating retinas or after retinal neuron damage (Chang et al., 2016; Yu et al., 2021). OPN is a secreted glycoprotein can act as a pro-inflammatory cytokine and is widely upregulated by various immune cells in response to injury across many tissues (Wang and Denhardt, 2008). Blocking OPN is neuroprotective against ganglion cell loss in a rat model of glaucoma (Yu et al., 2021). Accordingly, we investigated whether OPN stimulates the activation of NFκB-signaling in the retina. OPN acts through receptors including CD44 and integrins (Itgb3 and Itgav) (Helluin et al., 2000; Weber et al., 1996; Yokosaki et al., 2005), and can directly activate NFκB signaling via the IKK complex (Philip and Kundu, 2003). We find OPN *(Spp1)*upregulated specifically in reactive microglia/macrophage in NMDA-damaged retinas via scRNA-seq **(Fig. 6a)**. We find upregulation of *Spp1* in microglia occurs after the rapid upregulation of proinflammatory cytokines such as *Il1a, Il1b* and *Tnf* (**Fig. 6c**) (Todd et al., 2019). Complimentary to these findings, we detected expression of putative OPN receptors in MG. *Cd44* was rapidly upregulated (3-6hrs; activated MG clusters) and then downregulated (24-48hrs) in MG returning to rest **(Fig. 6b,d**). By comparison, levels of *Itgav* were downregulated, while the percentage of expressing cells was increased, in activated MG (6hrs after NMDA) and further downregulated in MG returning to resting **(Fig. 6b,d**). To determine if OPN influences NFκB-signaling in MG, we performed a single intravitreal injection of recombinant mouse OPN in NFκB-eGFP mice and harvested retinas 24h post-injection. We found induction of NFκB reporter in MG in undamaged retinas following OPN treatment **(Fig. 6e)**. Collectively, these findings suggest that OPN produced by reactive microglia/macrophage may act at *Cd44* and *Itgav* on MG to activate NFκB-signaling.

**Figure 6:**
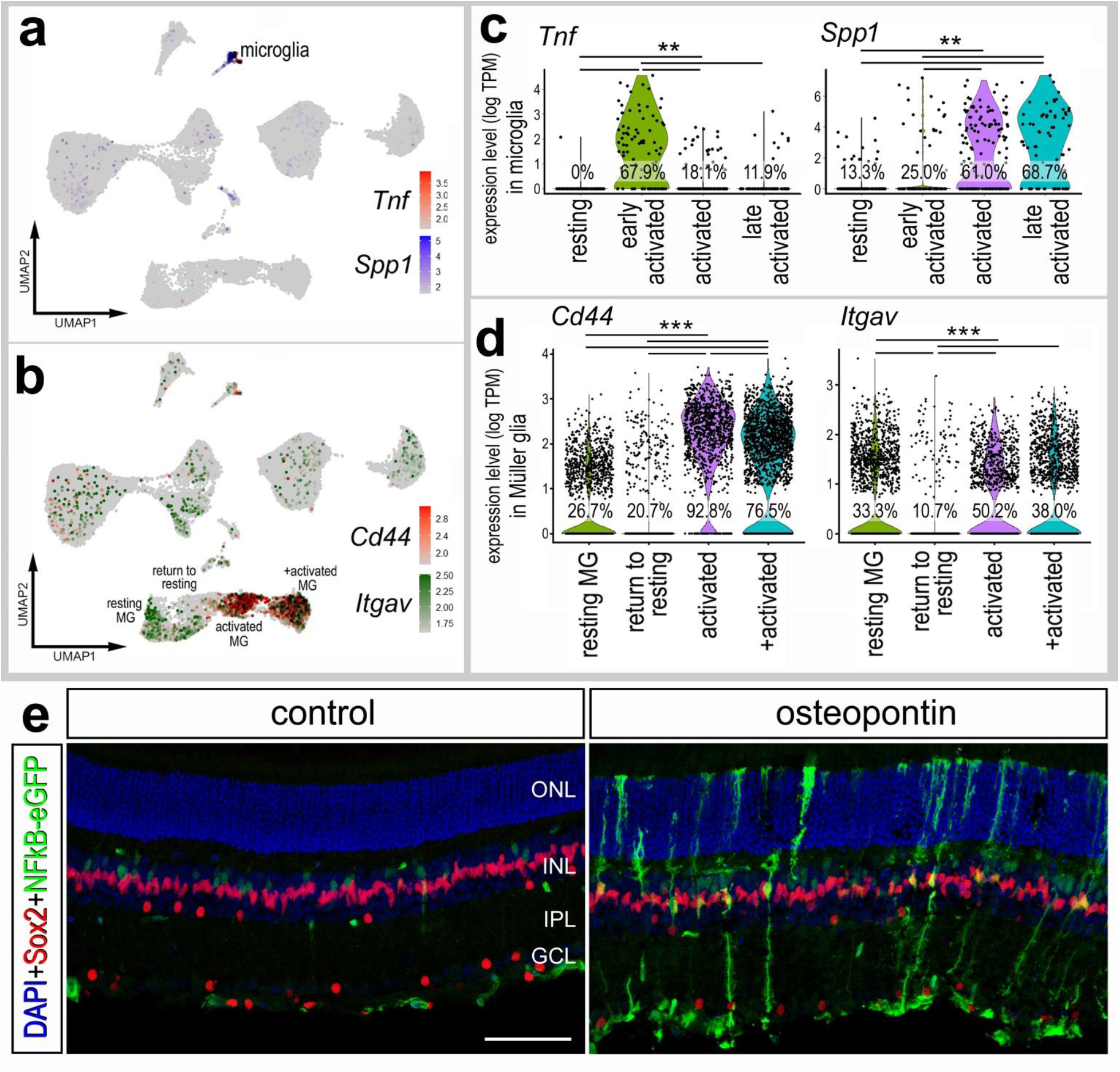
Osteopontin is expressed by microglia and activates NFκB signaling in MG. **(a-d)** Aggregated scRNA-seq libraries were prepared from WT control retinas and retinas 3h, 6h,12h, 24h, 36h, 48h, and 72h after NMDA damage. UMAP feature plots illustrate patterns of expression of *Tnf and Spp1* **(a)** or *Cd44* and *Itgav* **(b)** across all retinal cell clusters. Violin plots illustrate levels of expression of *Tnf* and *Spp1* in microglia (**c**), and *Cd44* and *Itgav* in MG (**d**). Each dot represents one cell. Significance (**p<10^-10^, ***p<10^-20^) of difference was determined by using a Wilcoxon Rank Sum with Bonferroni correction. (**e**) Eyes of NFκB-eGFP mice were injected with a single dose of recombinant mouse Osteopontin, retinas were harvested 24hr later, and retinal sections were labeled with DAPI (blue) and antibodies to GFP (green), Sox2 (red). Calibration bar represents 50 μm. Abbreviations: ONL – outer nuclear layer, INL – inner nuclear layer, IPL – inner plexiform layer, GCL – ganglion cell layer.

### Activation of NFκB-reporter in the optic nerve following crush

Lastly, we investigated whether injury other than excitotoxicity influenced NFκB reporter activity in the retina. We applied an optic nerve crush model, which damages axons and result in the death of retinal ganglion cells (Kalesnykas et al., 2012). We performed compression of the optic nerve, as described in methods, and harvested nerves and associated retinas 3 days post-injury (dpi), 5dpi, and 8dpi. We labeled corresponding retina’s with Brn3b to identify surviving retinal ganglion cells (RGCs). ONC resulted in a significant (p<0.0001, n=6) decrease (85.5 ± 12.7%) in the number of surviving Brn3b+ RGCs by 5 days after treatment (**Fig. 7a**). We did not detect activation of the NFκB-reporter within the retina at any time following ONC (**Fig. 7a,b**). However, in some eyes we observed a few NFκB-eGFP-positive MG in close proximity to the optic nerve head following ONC (**Fig. 7g, g’**). In undamaged optic nerves and optic nerve heads, we observed very few cells expressing the NFκB-reporter (**Fig. 7c,c’,e,f**). By comparison, we observed an increase in the number of cells expressing the NFκB-reporter throughout the optic nerve and within the optic nerve head at 5dpi and 8dpi (**Fig. 7d, d’, e, *e’*, g**). Many of the eGFP-positive cells within damaged optic nerves and optic nerve heads were co-labeled for Sox2 (**Fig. 7e,e’,g,g’**), a transcription factor known to be expressed by glial cells within the optic nerve and nerve head of many mammals including mice (Fischer et al., 2010).

**Figure 7:**
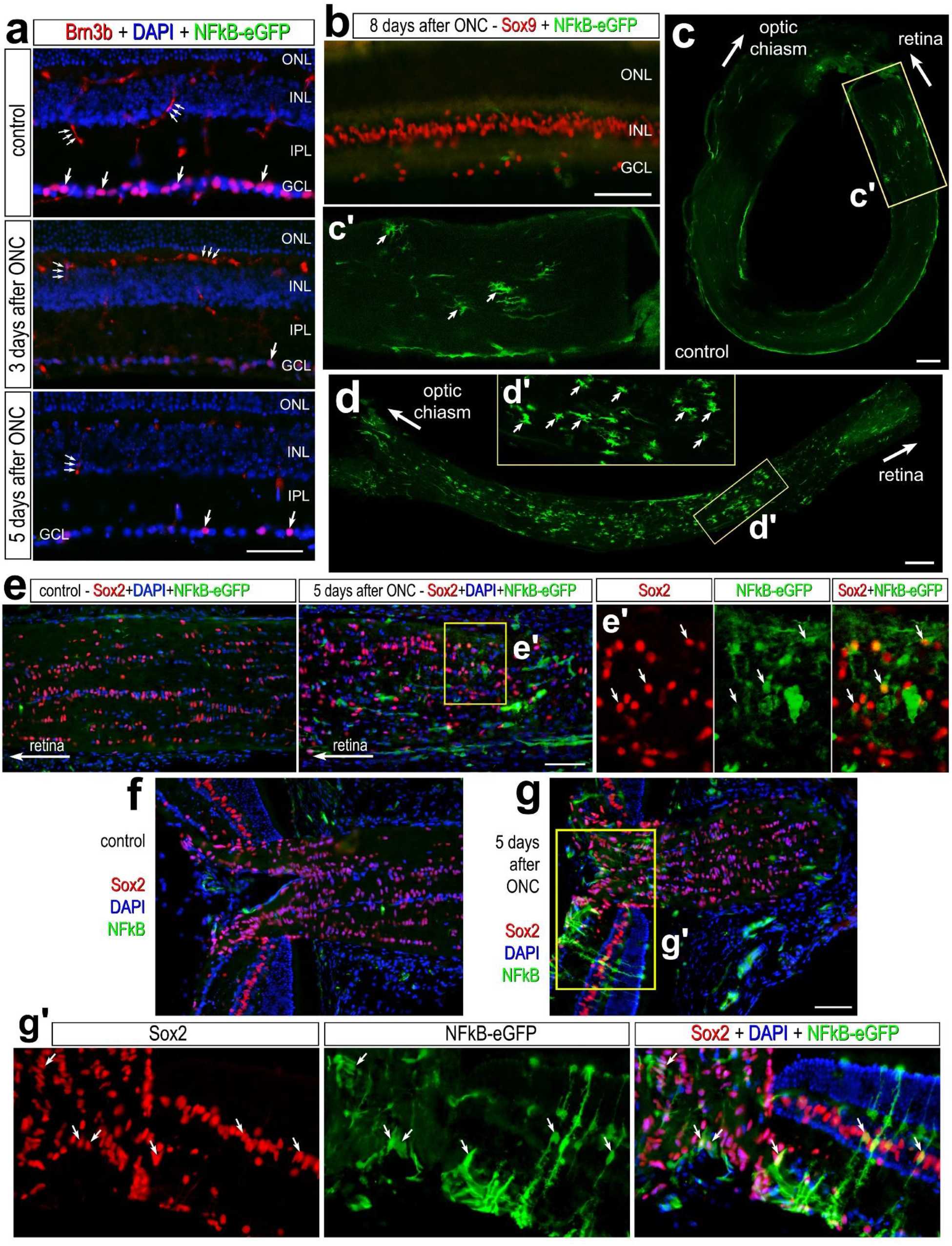
NFκB activation in glial cells following optic nerve crush. Optic nerves were crushed via forceps compression as described in methods, and retinas and optic nerves were harvested 3 days post-injury (dpi), 5dpi, and 8dpi. **(a)** Retinal sections of the were labeled with antibodies to Brn3b (red) and GFP (green). **(b)** Retinal sections 8dpi were labeled for Sox9 (red) and GFP (green). Whole-mounts of un-damaged **(c)** and crushed **(d)** optic nerves were labeled with antibodies to GFP. **c’** and **d’** are enlarged regions outlined within **c** and **d**. Large arrows indicate the direction of the optic nerve adjacent to the retina or towards the optic chiasm, and small arrows indicate GFP^+^ cells. Calibration bars represent 50 μm. **(e)** Longitudinal sections of control and damaged optic nerves harvested 5dpi were labeled for GFP (green) and Sox2 (red). **e’** is an enlarged region outlined in **e. (f-g)** Section of optic nerve head from control **(f)** and damaged **(g)** eyes 5dpi labeled for Sox2 (red) and GFP (green). **g’** is an enlarged region within **g**. Abbreviations: ONL – outer nuclear layer, INL – inner nuclear layer, GCL – ganglion cell layer.

## Discussion

Inflammatory signaling acts differentially across species to influence the regenerative capacity of MG after damage. It is important to understand how pro-inflammatory signaling fits into the complex network of pathways that control MG reprogramming. NFκB signaling is a master regulator of pro-inflammatory cascades and is differentially activated in MG across species, wherein expression is not highly induced in zebrafish MG but is expressed in mouse MG in response to retinal damage (Hoang et al., 2020; Palazzo et al., 2022). In the zebrafish, proinflammatory cytokines including TNF and IL6 are required to initiate the process of retinal regeneration from MG (Nelson et al., 2013). In the mouse retina, microglia repress the neurogenic reprogramming capacity of MG after damage (Todd et al., 2020), while microglia are necessary for initiation of MG reprogramming in the chick and fish (Fischer et al., 2014b; White et al., 2017). Consistent with this, we have shown that NFκB signaling is highly activated and suppresses neurogenic programs in mouse MG after damage (Palazzo et al., 2022). However, when microglia are ablated prior to damage, MG failed to activate NFκB signaling (Palazzo et al., 2022). Here we report that MG remained responsive to NFκB activation upon treatment with IL1β or TNF. Considering that microglia are the only cells that upregulate expression of IL1β and TNF after NMDA damage in the mouse retinas (Todd et al., 2019), we propose that NMDA-induced retinal damage results in microglial activation and secretion of proinflammatory cytokines, including IL1β and TNF, which stimulates NFκB-signaling in MG, and activates different pro-/anti-reprogramming related pathways to ultimately repress neurogenic reprogramming of MG in the mammalian retina (Fig. 8). This is consistent with other studies showing microglia derived TNF activates NFκB in mammalian MG *in vitro* (Ji et al., 2022), and microglia in the zebrafish retina upregulate TNF in response to NMDA damage (Iribarne and Hyde, 2021). However, in the zebrafish retina, TNF promotes MG reprogramming (Nelson et al., 2013), likely because NFκB gene modules are not highly upregulated in response to damage paradigms that stimulate microglia to produce TNF (Hoang et al., 2020).

**Figure 8:**
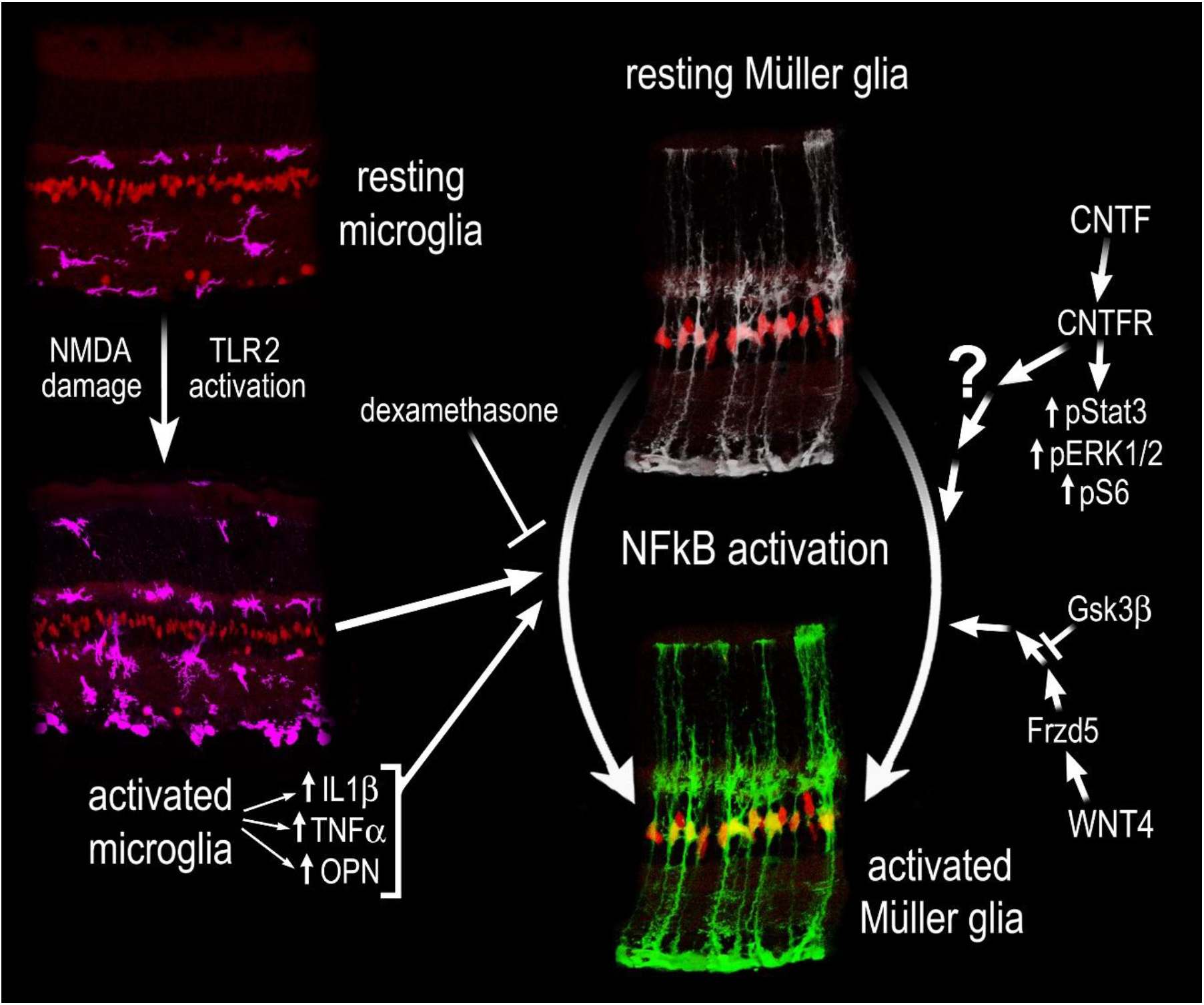
Schematic Summary of Results. Cropped images of retinal sections illustrating MG, labeled with Sox2 (red), resting and activated microglia labeled with Iba1 (magenta), and resting and activate MG (NFκB-GFP reporter - green). NMDA damage or TLR activation stimulates microglia to become reactive. Eeactive microglia secrete pro-inflammatory cytokines, including IL1β, TNFβ, and OPN, which act to stimulate NFκB signaling in MG. Additionally, WNT- and CNTF-signaling activate NFκB signaling in MG, but the nature of cross talk with Jak/Stat, mTOR, and MAPK signaling remains unclear.

We identified that activation of TLR receptors results in activation of NFκB in MG. Our data indicate that activation of NFκB in MG in response to treatment with TLR agonists is indirect and is mediated by microglia. We found TLR expression largely restricted to the microglia in the retina, and agonizing TLRs in microglia ablated retinas did not induce activation of NFκB in MG. TLRs are potently activated by HMGB1 (Park et al., 2004). Previously, we found that *Hmga/b/n* isoforms are highly upregulated in reactive MG after NMDA damage (Hoang et al., 2020) and inhibition or conditional knockout of NFκB signaling in MG diminishes expression of different *Hmg* isoforms (Palazzo et al., 2022). Thus, we hypothesize that there may be bi-directional communication between microglia and MG glia mediated by HMGB1. We hypothesize a potential mechanism by which MG secrete HMGB1, which then stimulate TLRs on microglia to induce pro-inflammatory cytokine production that subsequently stimulates NFκB-signaling in MG (Fig. 8). Inhibition of NFκB in MG breaks this feed-forward loop of driving inflammatory signaling between MG and microglia. Further studies into the role of HMGB1 are required to assess this hypothesis.

Jak/Stat signaling is involved in mediating cytokine signaling, and also promotes MG de-differentiation and proliferation of MG progeny in the fish and chick (Nelson et al., 2012; Todd et al., 2016). However, Jak/Stat signaling biases MG-derived progenitor cells towards a glial fate and represses neuronal differentiation in the chick (Todd et al., 2016). This is similar to our previous findings that NFκB signaling promotes pro-glial/anti-neuronal gene networks (Palazzo et al., 2022). NFκB likely converges on Stat3 in the retina, considering data showing that TNF activates Stat3 in the fish (Nelson et al., 2013). Here we find that CNTF treatment activates Jak/Stat and NFκB signaling (Fig. 8), but the nature of this crosstalk is still not well understood. CNTF receptor is expressed by MG and inner retinal neurons, but treatment with CNTF activates cell signaling only in MG. This suggests that CNTF-mediated signaling is (i) repressed in neurons, (ii) that there are deficits in signal transducers associated with CNTFR or (iii) or that cell signaling pathways other than those that we probed for were activated in neurons following treatment with CNTF. Further experiments are required to identify exactly how these pathways interact, converge and form a hierarchy to control glial responses to retinal damage.

There is evidence of both positive and negative bi-directional communication between Wnt- and NFκB-signaling, which is complex and context dependent (reviewed by (Ma and Hottiger, 2016)). In the chick retina, Wnt/β-catenin signaling is activated in response to NMDA damage, and inhibition of Wnt-signaling after damage attenuates MG-derived progenitor cell formation and proliferation (Gallina et al., 2015). Additionally, Wnt activation alone is sufficient to stimulate MG reprogramming in the absence of damage in the zebrafish retina (Ramachandran et al., 2011). Additionally, WNT/β-catenin-signaling promotes MG reprogramming in the mammalian retina (Osakada et al., 2007; Yao et al., 2016). Here we find that Wnt-signaling activates NFκB in the absence of damage, and our previous data indicates that NFκB signaling promotes pro-glial and anti-neurogenic programs (Palazzo et al., 2022). WNT4-mediated activation of NFκB signaling in our studies may be independent of β-catenin signaling. In addition to WNT4 treatment, we find inhibition of GSK3β is also sufficient to induce NFκB activation in MG. In the absence of WNT signaling, GSK3β is actively involved in the “destruction complex” that targets β-catenin for degradation. Upon Wnt ligands binding to Frizzled and LRP co-receptors, GSK3β is inactivated, allowing for accumulation and nuclear translocation of β-catenin (van Noort et al., 2002). There is evidence of crosstalk with

NFκB at multiple levels of this signaling cascade. Wnt/β-catenin activation elevates the ubiquitin ligase receptor (betaTrCP), which targets IκBα for degradation and consequently results in NFκB transactivation (Ma and Hottiger, 2016). It has been shown that overexpression of GSK3β results in repression of TNF induced NFκB activation and stabilizes the NFκB inhibitory IkB complex, thus preventing activation of NFκB signaling (Saegusa et al., 2007). Thus, it is likely that Wnt-mediated destabilization of GSK3β in MG is responsible for inducing NFκB activity independent of retinal damage (Fig. 8). We found that WNT receptors were predominantly expressed by MG, consistent with the selective activation of NFκB in these cells. However, our data indicate that GSK3β is expressed by most types of retinal glia and neurons, in addition to the MG. Thus, the mechanisms underlying the restricted activation of NFκB in MG in response to GSKβ inhibitors remains unclear.

We conclude that the NFκB pathway is dynamically activated in retinal MG in response to modulation of various cell signaling pathways. NFκB is activated predominantly in MG in response to excitotoxic damage of retinal neurons, whereas optic nerve crush activated NFκB in glial cells in the optic nerve, not in the retina. Activation of NFκB in MG depends on signals produced by reactive microglia, including proinflammatory cytokines IL1β and TNF. Broad inhibition of inflammation via dexamethasone effectively suppresses NFκB activation in MG, promotes neuronal survival, and suppresses the accumulation of immune cells in damaged retinas. The NFκB pathway can be activated via crosstalk from different cell signaling pathways including TLR-, CNTF/Jak/Stat-, Osteopontin, and Wnt/GSK3b-signaling, but not FGF2 (Fig. 8). We conclude that NFκB is a key cell signaling hub in retinal MG that is activated by different growth factors microglia-derived cytokines.

## Acknowledgements

We would like to acknowledge Dr. Dennis Guttridge for proving the NFkB reporter mouse line and Dr. Heithem El-Hodiri for his comments on the manuscript.

## Funding

This work was supported by RO1 EY022030-09 and R01 EY032141-02 (AJF).

## Notes

### Competing Interest Statement

The authors have declared no competing interest.

